# Probability cueing of singleton-distractor locations in visual search: priority-map‐ or dimension-based inhibition?

**DOI:** 10.1101/454140

**Authors:** Bei Zhang, Fredrik Allenmark, Heinrich R. Liesefeld, Zhuanghua Shi, Hermann J. Müller

**Affiliations:** General and Experimental Psychology, Department of Psychology, LMU Munich, Germany; Department of Psychological Science, Birkbeck College (University of London), London, UK

**Keywords:** search guidance, attentional capture, statistical (distractor location) learning, distractor suppression

## Abstract

Observers can learn the likely locations of salient distractors in visual search, reducing their potential to capture attention (Ferrante et al., 2018; Sauter et al., 2018a; Wang & Theeuwes, 2018a). While there is agreement that this involves positional suppression of the likely distractor location(s), it is contentious at which stage of search guidance the suppression operates: the supra-dimensional priority map or feature-contrast signals within the distractor dimension. On the latter account, advocated by Sauter et al., target processing should be unaffected by distractor suppression when the target is defined in a different (non-suppressed) dimension to the target. At odds with this, Wang and Theeuwes found strong suppression not only of the (color) distractor, but also of the (shape) target when it appeared at the likely distractor location. Adopting their paradigm, the present study ruled out that increased cross-trial inhibition of the single frequent (frequently inhibited) as compared to any of the rare (rarely inhibited) distractor locations is responsible for this target-location effect. However, a reduced likelihood of the target appearing at the frequent vs. a rare distractor location contributes to this effect: removing this negative bias abolished the cost to target processing with increasing practice, indicative of a transition from priority-map‐ to dimension-based – and thus a flexible locus of – distractor suppression.

**Public Significance Statement:** Distraction by a salient visual stimulus outside the ‘focus’ of the task at hand occurs frequently. The present study examined whether and how ‘knowledge’ of the likely location(s) where the distractors occur helps the observer to mitigate distraction. The results confirmed that observers can learn to suppress distracting stimuli at likely locations. Further, they showed that, the suppression may occur at different levels in the hierarchically organized visual system where the priorities of which objects to be attended in the environment are determined.

## INTRODUCTION

Recently, there has been a growing interest in statistical, location-probability learning in visual search. While most of this research has focused on the learning of target locations (e.g., Druker & Anderson, 2010; Geng & Behrmann, 2002, 2005; Walthew & Gilchrist, 2006; see also Miller, 1988; Müller & Findlay, 1987; Shaw & Shaw, 1977), more recently, there have been various attempts to extend this to the learning of distractor locations (e.g., Ferrante, Patacca, Di Caro, Della Libera, Santandrea, & Chelazzi, 2018; Goschy, Bakos, Müller, & Zehetleitner, 2014; Leber, Gwinn, Hong, & O’Toole, 2016; Sauter, Liesefeld, Zehetleitner, & Müller, 2018a; Wang & Theeuwes, 2018a). Collectively, these studies showed that observers appear to be able to learn, from experience, the spatial distribution of salient but task-irrelevant singleton or ‘pop-out’ distractors in the visual search array, to minimize the interference – or potential for ‘attentional capture’ – normally caused by such distractors. This appears to be the case whether the salient distractor occurs consistently at one specific, ‘most frequent’ location in relatively sparse displays (e.g., 4-item displays in Ferrante et al., 2018; 8-item displays in Wang & Theeuwes, 2018a, b) or within a ‘frequent’ region encompassing multiple possible locations, such as a whole display half, in dense displays (39-item displays in Goschy et al., 2014, and Sauter et al., 2018a). However, even though there is no dispute about the fact that observers can learn the statistical distribution of salient distractors and use this ‘knowledge’ to minimize the interference of distractors occurring at frequent locations, conclusions differ with regard to the locus, or processing stage, the functional architecture of search guidance at which the observed reduction of distractor interference (for frequent vs. infrequent locations) is realized.

Wang and Theeuwes (2018a), for instance, concluded that the interference reduction is inherently ‘spatial’ in nature. That is, in terms of the functional architecture of search guidance, learnt positional distractor suppression operates at the level of the ‘overall-saliency’ (Guided Search; e.g., Wolfe & Gancarz, 1997; Wolfe, 2007) or ‘priority’ (e.g., Fecteau & Munoz, 2006) map of the search array, essentially inhibiting any saliency signals at the frequent distractor location and thus preventing distractors at this location from summoning attention. This conclusion was based on one critical finding, namely, that not only distractor signals were suppressed at the frequent distractor location (as compared to distractors at infrequent locations), but also target signals. Of note in this context, Wang and Theeuwes (2018a) employed Theeuwes’ standard ‘additional-singleton’ paradigm (e.g., Theeuwes, 1992), in which the singleton target was shape-defined (an odd-one-out circle in a circular array of diamonds) and the distractor, when it appeared on a trial, was color-defined (the only red [diamond] shape amongst green shapes) – where the red singleton is typically more salient than the shape singleton, as evidenced by faster reaction times (RTs) when the target is a color singleton rather than a shape singleton. Thus, the fact that not only the distractor (color) singleton but also the target (shape) singleton was suppressed when it appeared at the frequent distractor location was taken as evidence that the suppression must operate on some superordinate or ‘global’ spatial representation that is essentially ‘feature-’ or ‘dimension-less’ in nature (as a result of which any singleton-feature item, whether color‐ or shape-defined, is effectively suppressed).

This evidence, and conclusion, is at variance with Sauter et al. (2018a). They found that, when the distractor and the target were defined in different visual dimensions (color-defined distractor and orientation‐ [i.e., in a sense, shape-] defined target, similar to Wang and Theeuwes (2018a), although after practice there was effective color distractor suppression in the frequent as compared to the rare distractor region, processing of the orientation-defined target was *not* impacted, that is: RTs to the target were *un*affected by whether the target appeared in the frequent distractor region or the rare region. However, this pattern changed in a condition in which the target and the distractor were defined within the same dimension, namely, orientation. In this case, while distractors appearing in the frequent region caused again less interference compared to distractors in the rare region (as in the different-dimension condition), targets were responded to slower when they appeared in the frequent (vs. the rare) distractor region. Sauter et al. (2018a) took this overall pattern to mean that, with different-dimension distractors, differential suppression of distractors in the frequent versus the rare region operates at a dimension-based level: stronger down-modulation of dimension‐ (in their study: color-) based feature contrast signals in the frequent versus the rare distractor region. In the same-dimension distractor condition, by contrast, down-modulation of distractor (orientation) signals would, as a consequence of intra-dimensional feature coupling (as assumed by the dimension-weighting account of Müller and colleagues; (e.g., Found & Müller, 1996; Müller, Heller, & Ziegler, 1995; Müller, Reimann, & Krummenacher, 2003), also down-modulate target (orientation) signals, thus impeding target processing while at the same time reducing distractor interference. However, down-modulating the distractor dimension might introduce a goal conflict, as the target is defined in the same dimension – which would require this dimension and coding of features within this dimension to be enhanced, rather than inhibited. Thus, instead, observers might resort to a global spatial suppression strategy – but in this case, too, not only distractor processing (beneficial) but also target processing (harmful) would be impaired in the frequent versus the rare distractor region. In a follow-up study, Sauter, Liesefeld, and Müller (2018b) provided evidence in favor of the latter alternative, namely, that, with same-dimension distractors, differential distractor suppression for the frequent vs. the rare region operates at a global spatial representation, that is: the overall-saliency or attentional-priority map.*^1^*

Thus, the pattern reported by Wang and Theeuwes (2018a; see also Wang & Theeuwes, 2018b) – characterized by impaired processing (leading to reduced interference) of color distractors at the frequent distractor location and at the same time impaired processing of shape targets at this location – is both empirically and theoretically at variance with Sauter et al. (2018a), according to whom such a pattern should be seen only with same-dimension distractors but not with different-dimension distractors. This raises the question why the different paradigms used by Wang and Theeuwes (2018a) and Sauter et al. (2018a) led to contradictory, if not diametrically opposite, data patterns and conclusions.

The present study, of two experiments, was designed to elucidate this question by examining (more or less subtle) differences of, as well as potentially special factors playing a role in, the Wang-and-Theeuwes vis à vis the Sauter-et-al. paradigm. Since the two experiments used variations of the Wang and Theeuwes (2018a) paradigm, this paradigm and some of the essential findings will be described in some detail in the next sections.

Adopting Theeuwes’ (1992) additional-singleton paradigm, observers were presented with a ring of 8 shape stimuli (radius: 4° of visual angle), one of which was designated a ‘target’ item: either the only circle in the array, presented amongst 7 diamond shapes; or the only diamond in the array, presented amongst 7 circular shapes (target-to-non-target assignment was changing randomly across trials). A target was present on all trials. The task was a ‘compound-search’ task: observers were required to find the target shape and respond to the orientation of a line within it, where a line of the same orientation (as in the target shape) or a different orientation appeared in each of the non-target shapes. All stimuli – except for possibly one: the additional singleton ‘distractor’ – were either green or red on a given trial. On distractor-present trials (67% of the total number), one non-target shape appeared in an odd-one-out color: either red (when the other items were green) or green (when the other items were red). The distractor could appear at any of the 8 possible locations, but was most likely to appear at one, ‘frequent’ distractor location (*p* = .65 on distractor-absent trials, as compared to *p* = .05 for each of 7 the remaining locations), randomly selected (and kept constant) for each observer. Although this was not expressly stated, it is clear from the analyses conducted that the distractor never coincided with the target location on distractor-present trials. On distractor-absent trials, the target appeared with equal likelihood at each location (including the frequent distractor location).

## EXPERIMENT 1

Experiment 1 was designed to replicate the findings by Wang and Theeuwes (2018a) and systematically examine whether their pattern of results can be attributed to *positional* inter-trial effects – which had not been (or, rather, could not be) examined by Wang and Theeuwes (2018a).

Note that in Wang and Theeuwes (2018a), a distractor occurred with 65% likelihood at the frequent distractor location, generating a substantial suppression effect for this location (and spreading to locations in the vicinity). In our own analyses (see Supplement in Sauter et al., 2018a; see also, e.g., Geyer, Müller, & Krummenacher, 2007; Kumada & Humphreys, 2002; Maljkovic & Nakayama, 1996), such inhibition effects do carry over across trials, that is: if a distractor on the current trial *n* falls at the same location as a distractor on the previous trial *n* − 1, then the interference effect caused by the current distractor is greatly reduced (owing to lingering inhibition placed on the ‘rejected’ distractor location on the previous trial – a type of cross-trial ‘inhibition-of-return’, IOR, effect); likewise, when the target on trial *n* appears at the same location as the distractor on trial *n* − 1, RTs to the target are increased. Now, when the distractor is (65%) likely to appear at one specific location, there are many positional inter-trial repetitions for this location (e.g., for two consecutive distractor-present trials, the likelihood of a repetition is .65×.65 = .4225 ; and for a distractor-absent trial following a distractor-present trial, the likelihood for a target falling at the previous frequent distractor location is .65×.125 = .0825). By contrast, there were 7 infrequent distractor locations, with a distractor appearing at each of these with a likelihood of 5% (i.e., .35/7). This made it very unlikely that two consecutive distractors appeared at the exact-same location (.05×.05 = .0025) and that, on a distractor-absent trial following a distractor-present trial, a target appeared at the same location as the distractor on the previous trial (.05×.125 = .00625).

Given these differential inter-trial contingencies, it is possible that at least some, if not all, of Wang and Theeuwes’s (2018a) critical effects – that is, the reduced distractor interference on distractor present trials for distractors at the frequent versus the infrequent locations, and the slowed RTs on distractor absent trials to targets at the frequent versus the infrequent locations – might be attributable to passive carry-over across trials of location-based inhibition, rather than to statistical learning of stimulus location probabilities. This would be consistent with Sauter et al. (2018a), who showed that when positional inter-trial ‘confounds’ are eliminated, there is no target location effect with same-dimension distractors.

Given that the number of trials in the Wang and Theeuwes (2018a) experiment (720 trials) was insufficient for such analyses, especially for estimating inter-trial effects on RTs to relatively rare cross-trial transitions, the number of trials in Experiment 1 was greatly increased to 3000 trials overall (administered in two separate sessions). This ensured some 29 observations, on average, per participant for the rare transitions with the target on trial *n* appearing at the exact same low-probability (distractor) location as a distractor on trial *n-1* (yielding a reasonably reliable measure of cross-trial inhibition for the rare distractor locations). In addition, it permitted us to examine for learning/practice effects in distractor suppression (e.g., Gaspelin & Luck, 2018; Geyer, Krummenacher, & Müller, 2008; Müller, Geyer, Zehetleitner, & Krummenacher, 2009; Zehetleitner, Goschy, & Müller, 2012; see also Cunningham & Egeth, 2016; Töllner; Conci, & Müller, 2015). In all other respects, Experiment 1 was identical in design and procedure to the study of Wang and Theeuwes (2018a).

### Method

#### Participants

A cohort of 24 participants (mean age: 28.33 years; age range: 18-40 years; 15 female) were recruited at Ludwig-Maximilians-University (LMU) Munich for this experiment. This sample size was determined based on the crucial target-location effect reported by Wang and Theeuwes (2018a). Although they did not report effect sizes, we calculated a *d*_z_ = 0.56 based on the reported *t* test. With α = .05, 1 – β = .80, and one-tailed testing (the direction of the effect was predicted: RTs to targets appearing at the inhibited, frequent-distractor location were predicted to be slowed, and not expedited!), the sample size needed to replicate this effect is 22 participants. As this is close to the 24 participants in the original Wang and Theeuwes study, we decided to collect the same number of participants, to be on the safe side. As we used a much larger number of trials, thereby reducing the measurement error in each individual average, we actually expected a much higher power. Indeed, post-hoc power calculations indicated a 1 – β = .9997 for Experiment 1 and 1 – β = .99 for session 1 of Experiment 2 (see below).

All participants were right-handed and all reported normal or corrected-to-normal vision, including normal color vision. They received 9 Euro per hour in compensation for their service. The study protocol was approved by the LMU Faculty of Pedagogics & Psychology Ethics Board. Informed consent was obtained from all participants prior to the experiment.

#### Apparatus

The experiment was conducted in a sound-reduced and moderately lit test room. Stimuli were presented on a CRT monitor at 1280×1024 pixels screen resolution and a refresh rate of 120 Hz. Stimuli were generated by Psychophysics Toolbox Version 3 (PTB-3) (Brainard, 1997) based on MATLAB R2016a (The MathWorks^®^ Inc). Participants viewed the monitor from a distance of 60 cm (eye to screen) and gave their responses by pressing the leftward‐ (‘horizontal) or upward-pointing (‘vertical’) arrow on the keyboard with their right-hand index or middle fingers, respectively.

#### Stimuli

The search displays (see Figure 1 for an example display) were composed of eight outline shapes (circles or diamonds) equidistantly arranged around a virtual circle with a radius of 4° of visual angle. The display items consisted of either one circle (target) and seven diamonds (non-targets), or, alternatively, one diamond (target) and seven circles (non-targets). The diameter of the circle shapes and, respectively, the side length of the diamond shapes was 2° of visual angle. Each outline shape contained a vertical or horizontal gray line inside (0.3° x 1.5°), with half of the internal lines being (randomly) vertical and half horizontal. In a certain percentage of trials (see below), one of the non-target shapes (the distractor) differed in color from all the other shapes, being either green (CIE [Yxy]: 22.5, 0.32, 0.55) amongst homogeneous red shapes (CIE [Yxy]: 8.82, 0.54, 0.36), or red amongst homogeneous green shapes. All search displays were presented on a black screen background (6 *cd*/*m*^2^), with a white fixation cross (1^°^ × 1^°^) in the center.

**Figure 1.**
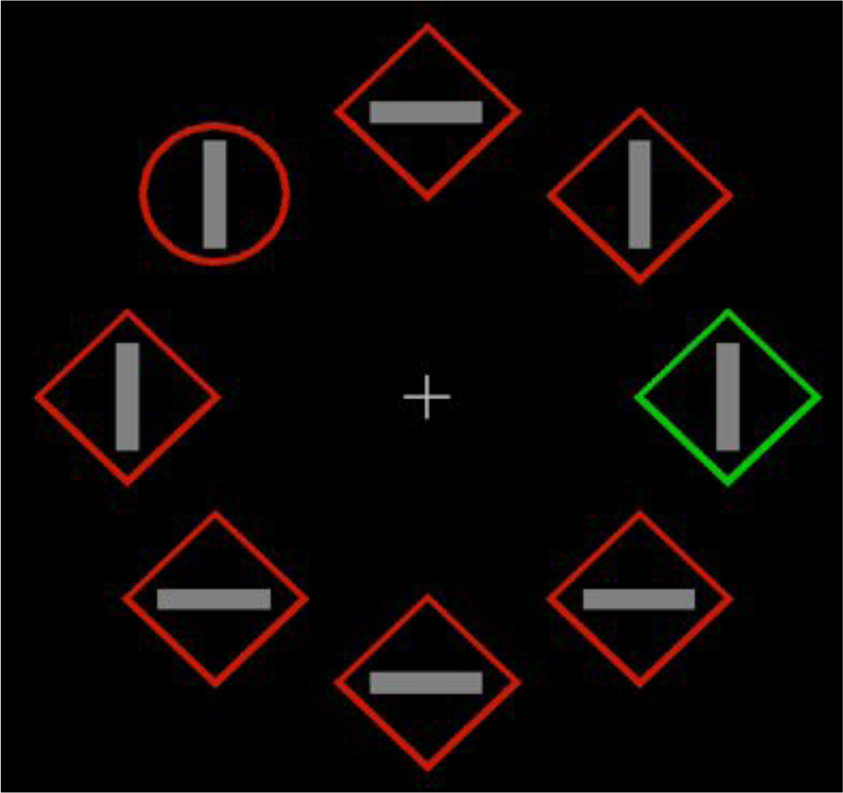
*Example of a visual search display. The search target is the singleton shape (here the only circle), and the distractor is a color singleton (here, the only green, diamond shape). Participants responded to the orientation of the bar inside the target shape (here vertical).*

#### Design

The target, which was present on all trials, was a singleton, odd-one-out shape amongst the 7 non-target shapes (either a circle or a diamond, randomly assigned on each trial). On trials without a distractor, the target was equally likely to appear at all 8 possible locations. On trials on which a distractor was present in the display, the target appeared equally frequently at all of the remaining 7 non-distractor locations. A singleton distractor, defined by a unique color (red or green, randomly assigned on each trial), appeared in 66% of the trials. If a distractor was present, it appeared with a likelihood of 65% at one, consistent location (frequent distractor location) and with a likelihood of 35%/7 at each of the other 7 locations (infrequent distractor locations). Note that the target and the distractor never appeared at the same location. The frequent distractor location remained the same for each participant, and was counterbalanced across participants. The experiment consisted of 3000 trials in total, subdivided into 2 sessions; each session was subdivided into 25 blocks of 60 trials each. Participants performed the two sessions on separate days.

#### Procedure

Each trial began with the presentation of a fixation cross for 500 ms, followed by the search array, which was shown until the participant gave a response. The intertrial interval (ITI) ranged between 500 and 750 ms (determined randomly). Participants were instructed to search for the target (the differently shaped item) and identify and respond to the orientation of the line inside – vertical or horizontal – as fast and as accurately as possible. For a vertical line, participants pressed the up arrow on the keyboard; and for a horizontal line the left arrow. At the end of the experiment, participants completed a post-experiment questionnaire, designed to determine whether they were aware of the frequent distractor location. This involved a two-stage procedure: first, participants had to indicate whether the distractor distribution was equal across all locations, or centered on one specific location; second, (even when they had given an equal response in stage 1) participants had to give a forced-choice response at which of the 8 locations the distractor had occurred most frequently (by marking the corresponding location on the ‘display’ depicted on the answer sheet). Prior to the main experiment (in each session), participants performed 60 unrecorded practice trials to re−/familiarize themselves with the task. Between trial blocks, participants could take a break of a self-determined length. Overall, each session took about one hour and 20 minutes to complete.

#### Bayes-Factor Analysis

Bayesian analyses of variance (ANOVAs) and associated post-hoc tests were carried out using JASP 0.9.0.1 (*http://www.jasp-stats.org*) with default settings. All Bayes factors for ANOVA main effects and interactions are ‘inclusion’ Bayes factors calculated across matched models. Inclusion Bayes factors compare models with a particular predictor to models that exclude that predictor. That is, they indicate the amount of change from prior inclusion odds (i.e., the ratio between the total prior probability for models including a predictor and the prior probability for models that do not include it) to posterior inclusion odds. Using inclusion Bayes factors calculated across matched models means that models that contain higher-order interactions involving the predictor of interest were excluded from the set of models on which the total prior and posterior odds were based. Inclusion Bayes factors provide a measure of the extent to which the data support inclusion of a factor in the model. Bayesian *t*-tests were performed using the ttestBF function of the R package ‘BayesFactor’ with the default setting (i.e., rscale =“medium”).

### Results and Discussion

All RT analyses below excluded outliers, defined as trials on which RTs were slower than 3 secs or faster than 150 ms (approximately 2% of trials, which is comparable to Wang and Theeuwes, 2018a), as well as trials on which participants made an incorrect response. In the analyses of inter-trial effects, the very first trial in each block was additionally excluded, because of the break between that trial and the last trial in the preceding block.

In the first instance, the data were analyzed analogously to Wang and Theeuwes (2018a), except that, since our experiment consisted of two sessions, we also examined for differences between the sessions reflecting practice effects. See Figures 2 (RTs and error rates as a function of target and distractor condition) and 3 (RTs and error rates as a function of the distance of the distractor from the frequent distractor location) for the results.

**Figure 2.**
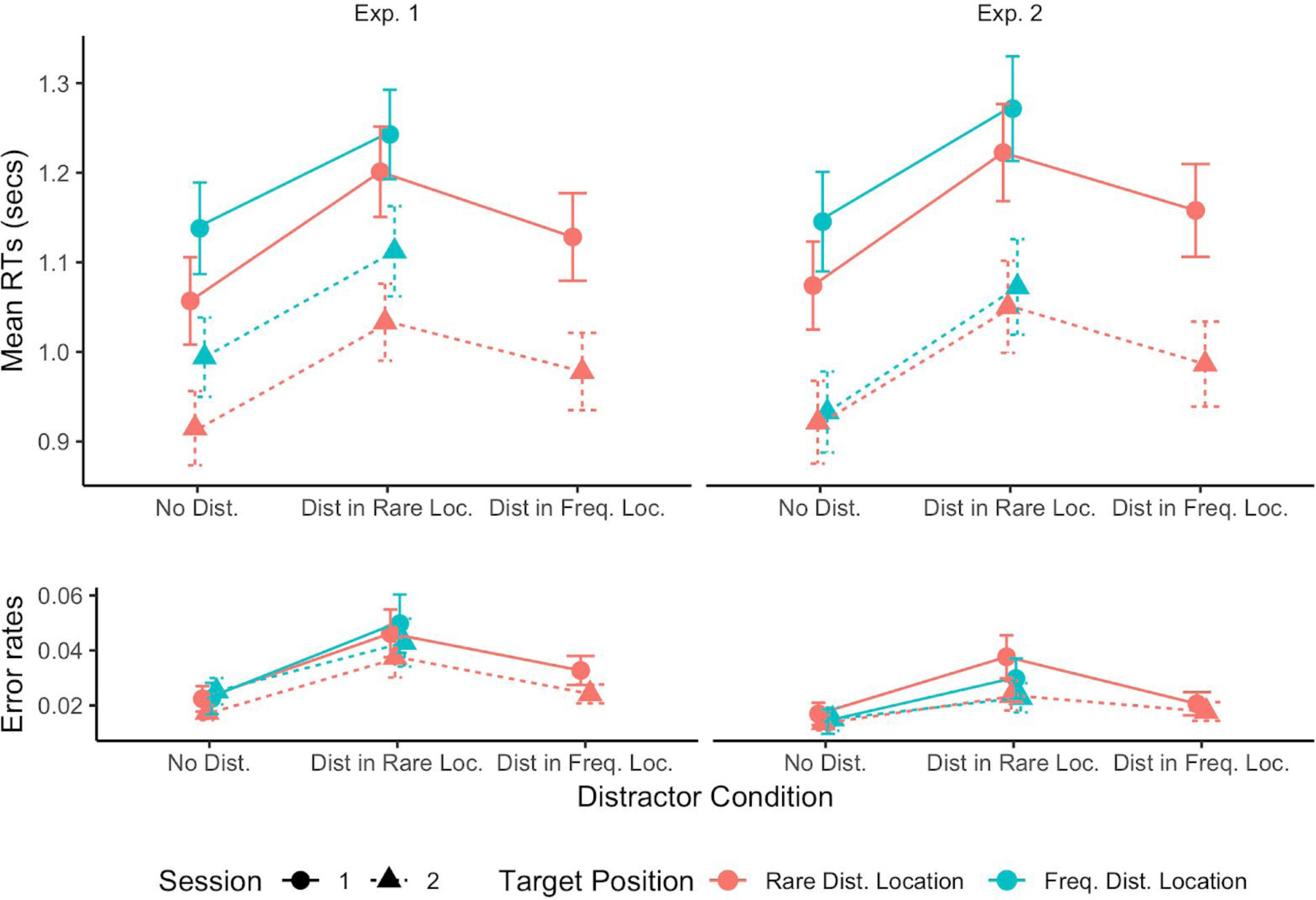
*Mean response times (RTs, upper panels) and mean error rates (lower panels) as a function of the distractor condition (distractor absent, at rare location, at frequent location), separately for the first and the second experimental session. Error bars denote one standard error. Note that the factor target position is defined only for distractor-absent trials and trials with a distractor at a rare location (on both of which the target could occur either at the frequent or at one of the rare distractor locations); on trials with a distractor at the frequent location, the target could appear only at one of the rare locations (as the target and distractor positions never coincided in the present paradigm).*

To examine how distractor presence at the high-frequency position compared to presence at one of the low-frequency positions affected RT performance, and whether the pattern differed between sessions, we performed a repeated-measures ANOVA with distractor condition (distractor absent, distractor at frequent location, distractor at rare location) and session (1, 2) as factors. Note that, similar to Wang and Theeuwes (2018a), this analysis disregarded the two target-position conditions (for distractor-absent trials and trials with a distractor at a rare location) depicted in Figure 2. This ANOVA revealed both main effects to be significant, distractor condition (*F* (46, 2) = 126.9, *p* < 0.001, 
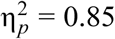
, *BF* > 100) and session (*F* (23, 1) = 37.8, *p* < 0.001, 
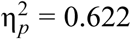
, *BF* > 100); the interaction was non-significant (*F* (46, 2) = 2.83, *p* = 0.07, *BF* = 0.14). The main effect of session reflected faster RTs in the second compared to the first session (mean RTs: 974 ms vs. 1123 ms). Concerning the main effect of distractor condition, post-hoc *t*-tests revealed that, relative to the distractor-absent baseline, there was significant RT interference wherever the distractor occurred (frequent distractor location, *t*(23) = 7.24, *p* < 0.001, *BF* > 100 ; rare locations, *t*(23) = 14.56, *p* < 0.001, *BF* > 100), but the interference was substantially reduced when the distractor occurred at the frequent location (1050 ms) compared to a rare location (1130 ms), *t*(23) = 9.53, *p* < 0.001, *BF* > 100 . The error rates (which were low overall: 3% on average) mirrored the RT pattern, effectively ruling out the observed RT effects were driven by differential speed-accuracy trade-offs.

Further, RTs to the target increased as the distractor on a given trial was presented further away from the frequent distractor location: an ANOVA with the factors ‘distance of distractor from frequent distractor location’ (ranging from dist-0 to dist-4) and session revealed the main effect of distance to be significant, (*F* (92, 4) = 19.6, *p* < 0.001, 
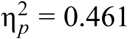
, *BF* > 100), without interacting with session (*F* (92, 4) = 0.38, *p* = 0.82, *BF* = 0.034). Importantly, the main effect of distance remained significant when dist-0, that is, the frequent location itself, was removed from the analysis (*F* (69, 3) = 6.20, *p* < 0.001, 
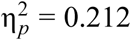
, *BF* = 1.21), indicating that distractor interference increased with distance of the distractor from the frequent location, rather than merely differing between the frequent and any other location. In particular, when the distractor was located adjacent to the frequent distractor location (dist-1), the interference effect (99 ms) was larger compared to dist-0 (57 ms) but smaller compared to greater differences (e.g., 141 ms for dist-2, dist-3, and dist-4 combined, which showed little difference amongst each other). This pattern was again mirrored in the error rates. Thus, a distractor appearing in close proximity to the frequent distractor location produced less interference than a distractor further away, consistent with a gradient of inhibition centered on the frequent distractor location.

Overall, these effect patterns replicate those reported by Wang and Theeuwes (2018a).

#### Distractor-absent trials

On trials without a distractor, responding to the target was significantly slower, by some 70 ms, when it appeared at the frequent distractor location compared to a rare location (see Figure 4), *t*(23) = 5.79, *p*<0.001, *BF*>100. Figure 4 depicts the RTs (on distractor-absent trials) as a function of the distance between the target location and the frequent distractor location. Although there was a significant effect of distance (*F*(92,4) = 12.56, *p*<0.001, 
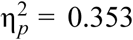
, *BF*>100), which did not differ between sessions (*F*(92,4) = 0.63, *p*=0.64, *BF*=0.043), there was little evidence of a gradient effect: while RTs were slower for dist-0, they differed little between the larger distances; there was actually no significant effect of distance after removing dist-0 (*F*(69,3) = 0.21, *p*=0.89, *BF*=0.029), and the RTs for dist-1 and dist-4 were virtually the same: 988 ms and 990 ms, respectively. – Again, the slowing of RTs to targets at the frequent distractor location replicate the effect reported by Wang and Theeuwes (2018a). As Wang and Theeuwes (2018a) did not report a distance analysis for distractor-absent trials, we cannot tell whether there was a significant gradient effect in their experiment. In any case, for distractor-present trials (for which Wang and Theeuwes reported a distance effect), based on Bayesian statistics, the evidence for a distance effect in Experiment 1 was also only weak when dist-0 was removed: *BF* = 1.21.

**Figure 4.**
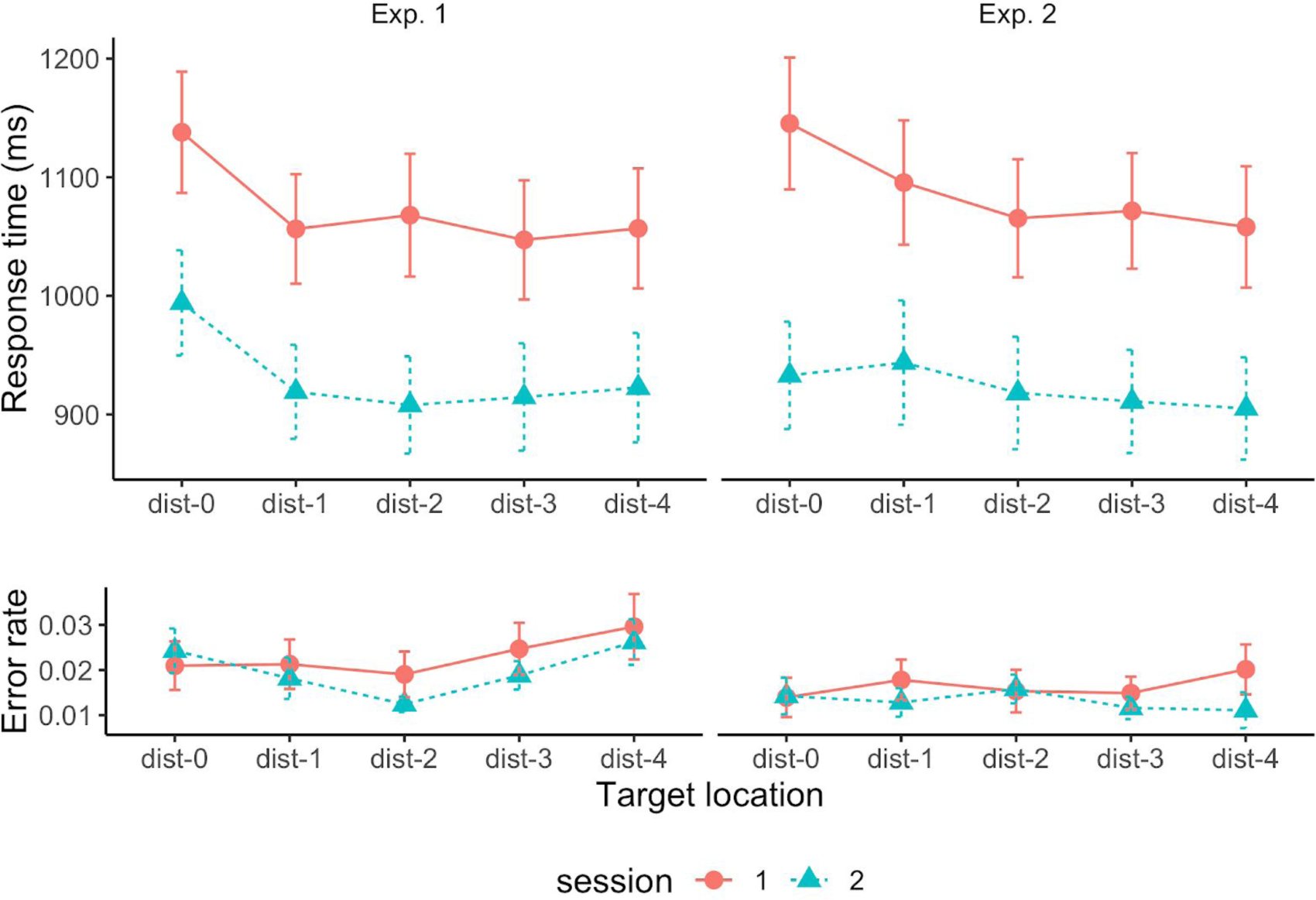
*Mean response times (RTs, upper panels) and mean error rates as a function of the distance of the target position from the frequent distractor position on distractor-absent trials, separately for the first and the second experimental session. Error bars denote one standard error.*

Next, going beyond Wang and Theeuwes (2018a), we examined for positional inter-trial effects along the lines outlined in the introduction to Experiment 1, in particular, we examined the effect of the distractor-to-target transition (same vs. different location) on trials *n-1* and *n* (distractor-absent trials *n*, with a distractor on trial *n-1*). This analysis focused on distractor-absent trials *n*, as this would be the condition that would reveal any carry-over (into trial *n*) of inter-trial inhibition of the distractor location on trial *n-1* in its purest form.

With a distractor absent on a given trial *n*, the target on this trial could appear either at the frequent distractor location or at one of the rare locations. As regards the distractor condition on the previous trial *n-1*, there are then three possibilities: the target and distractor locations are either coincident (i.e., target *n* appears at the same location as the distractor on trial *n-1*) or non-coincident, that is, target *n* appears at a location different to that of the distractor on trial *n-1* or there was no distractor on trial *n-1* (i.e., there were two consecutive distractor-absent trials). As the latter two conditions revealed little difference, we collapsed them into one, ‘non-coincident’ condition. Figure 5 shows how RTs and error rates depend on target-distractor coincidence for each target condition (target at frequent, rare distractor location) and session. Overall, there appeared to be some effect of target-distractor coincidence – indicative of cross-trial inhibition – for targets on trial *n* appearing at the location of a rare distractor on trial *n-1* (1013 vs. 985 ms, *t*(23) = 1.85, one-tailed*^2^ p* = 0.039, *BF* = 0.93), but there was no effect whatsoever for targets appearing at the frequent distractor location (1061 vs. 1069 ms, *t*(23) = 0.52, one-tailed *p* = 0.70, *BF* = 0.24). The effect for rare distractor locations appeared to be driven mainly by the second session (second session, coincident vs. non-coincident: 976 vs. 913 ms; *t*(23) = 3.15, *p* < 0.05, *BF* = 9.5): an ANOVA with the factors target condition (target at frequent, rare distractor location), target-distractor coincidence, and session suggested the pattern of RTs as a function of target condition and target-distractor coincidence to differ across sessions (three-way interaction: *F* (23, 1) = 5.664, *p* = 0.026, 
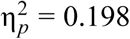
, *BF* = 0.57). However, this interaction is put into question by the Bayes factor. In any case, the (if anything) larger inter-trial inhibition associated with rare distractor locations is at variance with the hypothesis that the strong overall-inhibition of the frequent distractor location arises as a result of stronger positional (inhibitory) cross-trial dynamics for the frequent location.

**Figure 5.**
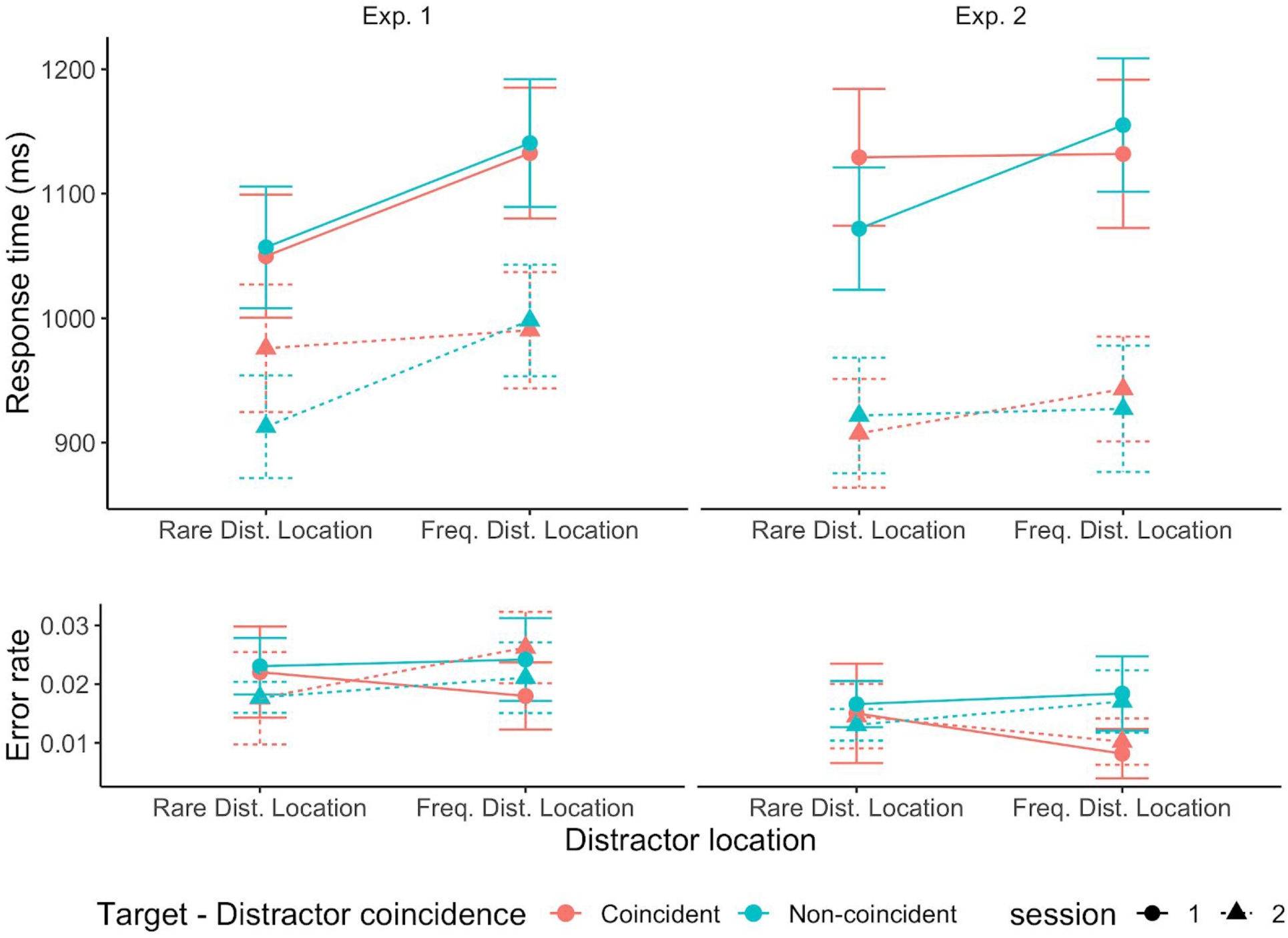
*Mean response times (RTs, upper panels) and mean error rates (lower panels) as a function of the target position (at the frequent distractor location, at a rare distractor location) and coincidence/non-coincidence of the target position with the previous distractor position, separately for the first and the second experimental session.*

#### Color-repetition effects

Because it is conceivable that participants attempt to suppress the distractor based on its color as well as its position, even though the target (i.e., non-distractor) and distractor color changed randomly from trial to trial, we also examined for an effect of repeating versus switching the color assignment between trials. Like Wang and Theeuwes (2018a), we first examined whether the amount of interference caused by a distractor at the frequent distractor location was different when color assignment was repeated compared to when it changed. Contrary to Wang and Theeuwes (2018a), we found the interference effect to be significantly reduced when the color assignment was repeated compared to when it changed (46 ms vs. 70 ms, *t*(23)=-3.19, *p*=0.0041, *BF*=10.4). This color-repetition benefit is indicative of some additional, color-feature-specific component of distractor suppression.

Given this finding, we went on to perform a more detailed analysis of the color-repetition benefits, more precisely: of the color repetition benefit as a function of the distractor condition (distractor absent, at rare location, at frequent location) on the current trial *n*, dependent on the distractor condition of *trial n-1*. As this analysis is exploratory and somewhat tangential to the question at issue in the present study, the results are detailed in a *Supplementary* section. In brief, this analysis revealed a color-repetition benefit on the current trial *n* only when a distractor appeared at one of the rare locations on the preceding trial *n-1* (not when there was no distractor or when a distractor appeared at the frequent location), and a benefit was evident both when the current distractor appeared at a rare location and when it appeared at the frequent location (but not when there was no distractor on the current trial). – This pattern is consistent with the idea that when a distractor at a rare location captures attention (which is more likely to occur in comparison with a distractor at the frequent, i.e., ‘spatially’ suppressed, location), the distractor color is inhibited in order to disengage attention from the rare distractor and re-allocate it to the target. If this color set (inhibition of the distractor color) is carried over across trials, it would diminish the potential of a distractor defined by the same color, wherever it appears in the display, to attract attention.

#### Summary

Thus, overall, our results provide a near-perfect replication of those reported by Wang and Theeuwes (2018a). In particular, there was a significant target location effect (on distractor-absent trials), with targets being responded to slower when they appeared at the frequent distractor location compared to one of the infrequent locations. Our analyses of positional intertrial effects revealed that, while there was evidence of cross-trial inhibition (IOR) for the rare distractor locations, there was no evidence of such an effect whatsoever for the frequent location. This pattern is at variance with an account of the target position effect in terms of asymmetric carry-over of inhibition (IOR) across trials between the frequent and rare distractor locations, and it is entirely in line with the interpretation put forward by Wang and Theeuwes (2018a), namely, that there is strong positional inhibition of the frequent distractor location. In fact, at least judging from the distractor-absent trials (on trial *n*), inhibition appeared to be saturated for this location, in that it was so strong that it is entirely unaffected by positional inter-trial effects! Also, there was no evidence that this pattern changed as a result of practice on the task: cross-trial inhibition was essentially absent for the frequent distractor location in both sessions/halves of the experiment (whereas it increased from session 1 to session 2 for the rare locations).

Going beyond Wang and Theeuwes (2018a), we found a significant benefit of repeating (vs. switching) the color of the distractor (relative to that of the other display items) across consecutive trials, though only when there was a distractor at a rare location (not when there was one at the frequent location) on the previous trial. This points to an element of color-based suppression of distractors at the frequent location, on top of space-based suppression. However, as color-based suppression works equally for all (potential distractor) locations (i.e., both the frequent and the 7 rare locations), this component cannot account for the overall reduced interference with distractors at the frequent versus the rare locations.

## EXPERIMENT 2

Experiment 1 showed that the result pattern of Wang and Theeuwes (2018a) cannot be reduced to positional inter-trial effects. Nevertheless, it still remains a question whether the (near-saturated) suppression of the frequent distractor location can be attributed solely to distractor position learning, that is, learning to ignore the frequent distractor location. This does remain an open question because, in the paradigm of Wang and Theeuwes, not only was a distractor more likely to appear at the frequent distractor location (on 65% of the distractor-present trials), but a target was also, at the same time, less likely to appear at this location. In number terms: on distractor-present trials, while a target appeared with a likelihood of 95%/7 (= 65%/7 + 30%/7) = approx. 14% at an infrequent distractor location, it appeared only with a likelihood of 35%/7 = 5% at the frequent distractor location; in other words, it was almost three times less likely to appear at the frequent distractor location. Accordingly, learning of the likely distractor location is potentially ‘confounded’ with learning of an unlikely target location, so that we cannot tell whether the suppression effect is due to one or the other or a combination of both. Experiment 2 was designed to examine for this, by making the frequent distractor location as likely to contain a target as any of the infrequent locations not only on distractor-absent trials, but also on distractor-present trials.

### Method

Methodologically, Experiment 2 was essentially the same as Experiment 1, the only exception being that, on distractor-present trials, a target was equally likely to appear at the frequent distractor location as at any one of the infrequent locations, by increasing the likelihood of a target appearing at the frequent distractor location on the 35% of trials on which a distractor occurred at an infrequent location. On distractor-absent trials, the target appeared equally likely at all locations, in any case. 24 new volunteers (mean age: 24.96 years; age range: 19-34 years; 16 female) participated in Experiment 2, on the same terms and procedural conditions as in Experiment 1. Overall, participants performed 3000 trials in two sessions, which again allowed us to examine for any changes in performance as a function of practice (session effects).

### Results and Discussion

Analogously to Experiment 1, we first examined the RTs (and error rates) by a repeated-measures ANOVA with distractor condition (distractor absent, at frequent location, at rare location) and session as factors. See the right-hand side of Figure 2 for a depiction of the results. This ANOVA revealed both main effects to be significant: distractor condition (*F* (34.2, 1.5) = 122.6, *p* < 0.001, 
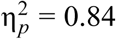
, *BF* > 100), and session (*F* (23, 1) = 45.2, *p* < 0.001, 
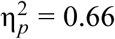
, *BF* > 100); the interaction failed to reach significance (*F* (46, 2) = 2.83, *p* = 0.07, *BF* = 0.13).

The session effect again reflected faster RTs in the second compared to the first session (mean: 973 ms vs. 1146 ms). Concerning the effect of distractor condition, post-hoc t-tests revealed that relative to the distractor-absent baseline, there was significant RT interference wherever the distractor occurred (frequent distractor location, *t*(23) = 11.99, *p* < 0.001, *BF* > 100 ; rare locations, *t*(23) = 12.98, *p* < 0.001, *BF* > 100), but the interference was significantly reduced when the distractor occurred at the frequent location (1070 ms) compared to the rare location (1150 ms), *t*(23) = 7.63, *p* < 0.001, *BF* > 100 . The error rates (which were low overall: 2% on average) mirrored the RT pattern, effectively ruling out that the observed RT effects were merely driven by speed-accuracy trade-offs. Essentially, this replicates the pattern seen in Experiment 1.

However, different to Experiment 1, in Experiment 2 there was no evidence of increased distractor interference with distance of the current distractor from the frequent location (see right panel of Figure 3): an ANOVA with the factors ‘distance of distractor from frequent distractor location’ (ranging from dist-1 to dist-4) and session revealed neither a significant main effect of distance nor a significant interaction with session (main effect:*BF* = 0.034 ; interaction: *F* (69, 3) = 0.085, *p* = 0.97, *BF* = 0.058). *F* (69, 3) = 1.011, *p* = 0.39,

**Figure 3.**
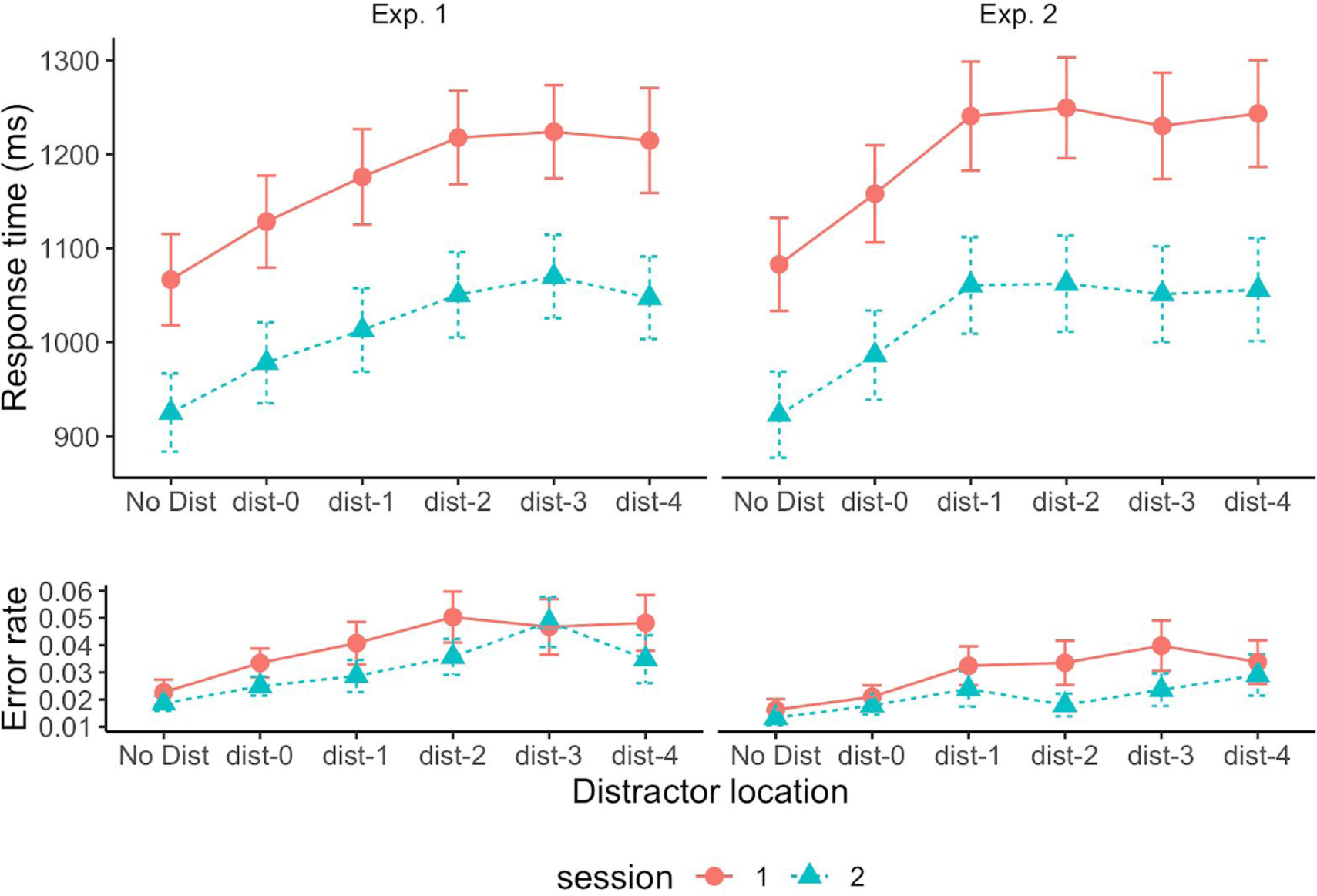
*Mean response times (RTs, upper panels) and mean error rates (lower panels) as a function of the distance of the distractor on a given trial from frequent distractor location, separately for the first and the second experimental session. Error bars denote one standard error. (dist-0 denotes distractor at frequent location, dist-1 denotes distractor adjacent to frequent location, etc.)*

To determine whether, on distractor-absent trials, there is any effect of target condition (at frequent distractor location vs. rare distractor location) after accounting for carry-over effects from distractor inhibition on the previous trial, we analyzed the effect of target condition after removing trials on which the target appeared in the previous distractor position (i.e., we considered the non-coincident condition; see right panel of Figure 5) in an ANOVA with target condition and session as factors. Unlike in Experiment 1, the effect of target condition differed between sessions (*F* (23, 1) = 6.11, *p* < 0.01, 
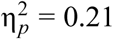
, *BF* = 1.71). In session 1, RTs were slower when the target appeared at the frequent distractor location compared to any other location (1141 ms vs. 1057 ms; *t*(23)=3.50, p<0.01, BF=19.6) – which mirrors the pattern seen in Experiment 1 and in Wang and Theeuwes (2018a). In session 2, by contrast, the difference was not statistically significant (927 ms vs. 922 ms; *t*(23) = 0.36, *p* = 0.72, *BF* = 0.23) – that is, there was no longer a target-location effect – a pattern consistent with Sauter et al. (2018a). – Like in Experiment 1, there was no evidence of a graded effect of the distance of the target location from the frequent distractor location (see right-hand side of Figure 4), not even in session 1, where there was a significant target-location effect (an ANOVA including only session 1 and removing dist-0 yielded no significant effect of distance: *F*(69,3) = 1.48, *p*=0.23, *BF*=0.27).

The pattern of positional inter-trial effects (carry-over of inhibition of the distractor location on distractor-present trial *n-1* to distractor-absent trial *n*) was overall similar to that seen in Experiment 1 (see right-hand side of Figure 5): collapsed across the two sessions, there was evidence of a carry-over of inhibition (RT coincident > RT non-coincident) for the rare locations (22-ms inhibition), but not the frequent location (3-ms difference in the opposite direction to inhibition). However, there was no significant interaction between coincidence and location (interaction coincident/non-coincident x target at frequent/rare location, *F*(23,1) = 1.26, *p*=0.27). This time, though, the effect appeared to be arising in the first session (*F*(23,1) = 3.48, *p*=0.075, 
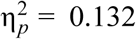
, *BF*=1.75; the three-way, session × coincidence × location, interaction was significant: *F*(23,1) = 7.36, *p*=0.012, 
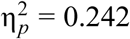
, *BF* = 1.08); in the second session, the frequent and rare locations appeared equally (un−)affected by positional cross-trial inhibition. However, looked at in terms of the Bayes factor, the evidence for an interaction involving the factor session is not convincing.

#### Color-repetition effects

As for Experiment 1, we first examined whether the amount of interference caused by a distractor at the frequent distractor location differed depending on the repetition versus change of the color assignment across consecutive trials. Again, and contrary to Wang and Theeuwes (2018a), there was a significant color-repetition (vs. −change) benefit (78 ms vs. 60 ms, *t*(23)=2.57, *p*=0.017, *BF*=3.1). A follow-up analysis of the color repetition benefit as a function of the distractor condition (distractor absent, at rare location, at frequent location) on trial *n*, dependent on the distractor condition of *trial n-1* (see Supplementary Section for details), revealed a similar picture to that seen in Experiment 1: there was a (numerical) color-repetition benefit on the current trial only when a distractor appeared at one of the rare locations on the preceding trial, and a benefit was evident when the current distractor appeared at a rare location (significant) and when it appeared at the frequent location (numerical), but not when there was no distractor on the current trial). By and large, this is in line with the account sketched for Experiment 1: when a distractor appears at a rare location on trial *n-1* (in which case it is likely to summon attention), its color may be noted and suppressed (to aid re-allocation of attention to the target location); this inhibitory set is then carried over across trials and benefits performance when the color assignment is repeated (by down-modulating the color feature contrast of the distractor on trial *n*, making it less potent to capture attract attention).

#### Summary

Overall, Experiment 2 in many respects replicates the findings of Experiment 1: observers do come to learn, and apply strong inhibition to, the frequent distractor location. However, there appears to be a major shift between the two sessions in how this inhibition operates. In session 1 (as in the whole of Experiment 1), it involves a robust target location effect, that is: targets are responded to slower when they appear at the frequent compared to one of the rare distractor locations – consistent with inhibition being applied (to the frequent distractor location) at the level of the supra-dimensional ‘priority’ map. In session 2, by contrast, the target location effect is no longer evident (in fact, the Bayes factor, *BF* = 0.23, favors the null hypothesis of no target location effect!), and this is so despite the fact that the magnitude of the distractor location effect (i.e., the difference in interference between distractors at the frequent vs. the rare locations) is virtually unchanged (72 ms in session 2 vs. 82 ms in session 1; *BF* = 0.13 for the distractor condition × session interaction). The lack of a target location effect mirrors the results of Sauter et al. (2018) for conditions in which the distractor is defined in a different dimension to the target: it is inconsistent with spatial inhibition of the distractor location operating at the level of the priority map, but consistent with inhibition operating at a dimension-based level, such as the map of (color-) dimension-specific feature contrast signals (which are then integrated across dimensions in the search-guiding priority map).

We propose that this reflects an adaptive shift of the processing level at which distractor inhibition is applied, which is adaptive to the distractor and target location probabilities prevailing in particular task scenarios. Of note, the beginning of a shift can be discerned already within the first session (see Figure 6): splitting the first session of Experiment 2 in halves, the target-location effect on distractor-absent trials turns out smaller in the second compared to the first half (first vs. second half of session 1: 91 vs. 51 ms; *t*(23)=2.69, *p*=0.013, *BF*=3.85*^3^*; the 51-ms effect in the second half is significantly different from zero: *t*(23)=3.03, *p*=0.006, *BF*=7.51), without a corresponding decrease of the interference reduction for distractors at the frequent versus the rare locations (first vs. second half: 78 vs. 85 ms; *t*(23)=0.50, *p*=0.62, *BF*=0.24). That is, the transition from priority-map‐ to dimension-based suppression of the likely distractor location occurs more gradually, but may need some 1500-plus trials to be completed.

**Figure 6.**
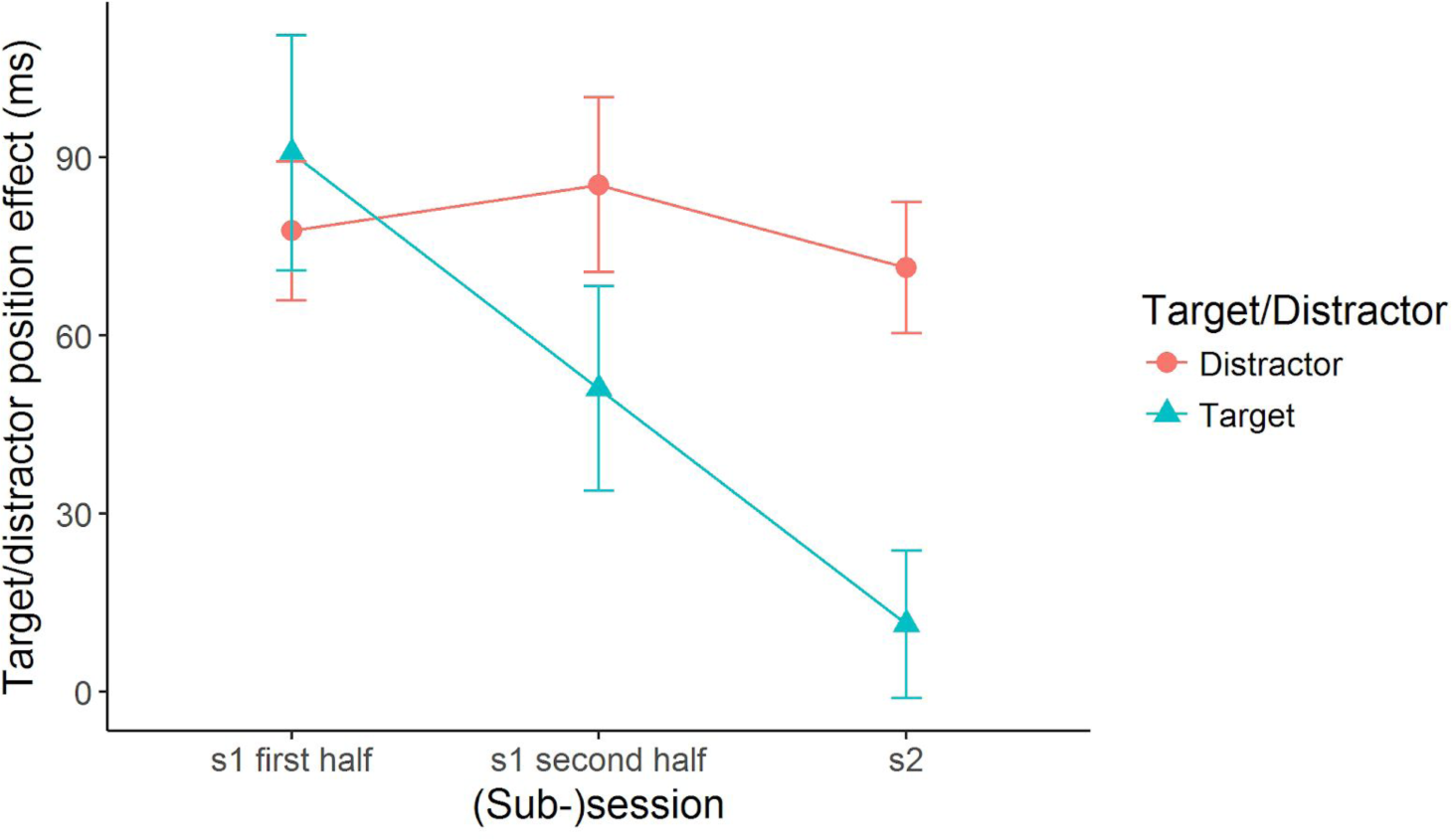
*Distractor-location effect (RT difference between conditions with a distractor at a rare vs. the frequent distractor location), and target-location effect (RT difference between conditions with the target at the frequent vs. a rare distractor location) on distractor-absent trials, across the course (first and second half of session 1, and whole of session 2) of Experiment 2. While the target-location effect reduces towards zero, the distractor-location effect remains virtually unchanged. (Data uncorrected for cross-trial inhibition of distractor locations.)*

## GENERAL DISCUSSION

Two experiments designed to examine the target-location effect in Wang and Theeuwes’ (2018a) paradigm revealed their pattern of effects to be highly replicable. In particular, in both experiments, there was strong suppression of the frequent distractor location: a distractor at this location caused substantially less interference than a distractor at a rare location (on distractor-present trials). In addition, in Experiment 1 (which was an exact replication of Wang and Theeuwes’s, 2018a, experiment, the only difference being an increased number of trials), we also found a target-location effect on distractor-absent trials: RTs were substantially slowed when the target appeared at the frequent distractor location compared to a rare location. Contrary to our initial hypothesis, however, this effect could not be reduced to the carry-over of positional inhibition of the distractor location from one (distractor-present) trial to the next (distractor-absent) trial being cumulatively stronger for the frequent (i.e., statistically frequently inhibited) distractor location compared to the rare locations. If anything, the effect pattern was the other way round: the frequent distractor location was inhibited (tonically) so strongly that it was virtually unaffected by cross-trial positional inhibition. This overall effect pattern was essentially the same in both experimental sessions.

Experiment 2 went on to examine whether this strong inhibition was influenced not only by the distractor-location probability, but also by the target-location probability. In Wang and Theeuwes’s (2018a) original paradigm, the frequent distractor location was actually some three times less likely to contain a target than any of the rare locations on distractor-present trials, effectively providing participants with a secondary incentive to ignore the frequent distractor location. This target-location bias was removed in Experiment 2, to examine the resulting effect pattern. The results revealed that in the first experimental session (averaged across the two session halves), the effect pattern essentially mirrored that obtained in Experiment 1. In particular, a distractor at the frequent (vs. one of the rare) location(s) caused less interference, and responding was significantly slowed when the target appeared at the frequent (vs. a rare) distractor location on distractor-absent trials. This pattern was changed in the second session: while distractor interference was still reduced – by, in fact, an equal amount – on distractor-present trials, there was no longer a target location effect (in fact, the Bayes factor argues in favor of a null effect). This is the very pattern observed by Sauter et al. (2018a) for conditions with a distractor defined in a different dimension (color in both studies) to the target (shape in the present study, orientation in the Sauter-et-al. study).

The effect pattern seen in Experiment 1 and session 1 of Experiment 2 is consistent with the notion, advocated by Wang and Theeuwes (2018a; see also Ferrante et al., 2018)^4^, that (spatial) suppression of the frequent distractor location operates at the level of the search-guiding priority map. This is beneficial in that is brings about a substantial reduction of distractor interference; at the same time, it is costly in that targets appearing at the frequent distractor location fall into the inhibitory trough: they take much longer to be detected and processed. By contrast, the effect pattern seen in session 2 (and already emerging during the second half of session 1) of Experiment 2 is consistent with the notion of dimension-based (spatial) inhibition, advocated by Sauter et al. (2018a): strongly inhibiting color signals at the frequent distractor location effectively reduced the interference of (color-defined) distractors at this location, while leaving the processing of shape/orientation-defined (target) signals unaffected. Thus, the present results argue that removal of the target location bias in Wang and Theeuwes’s (2018a) paradigm brings about an adaptive shift from priority-map-based suppression to dimension-based suppression.

There are at least two questions to be discussed as regards this interpretation: (i) why is the target-location effect fully abolished only in session 2 of Experiment 2, but not already in session 1 (even though it started to decrease in the second half of session 1)? (ii) is the mode of suppression applied (i.e., the level, in the functional architecture, at which suppression operates) flexible, a matter of strategic set?

Concerning question 1, one plausible answer is that observers first pick up the more striking distractor location probability (as also evidenced by the fact that most observers became consciously aware of the likely distractor location*^5^*), and this makes them operate a purely spatial, priority-map-based inhibitory strategy: suppress any stimulus at this location, because it is likely to be a distractor. However, over time, they come to realize that this strategy harms detection of (and responding to) the target when it appears at the frequent distractor location, especially when they come to learn more slowly, in Experiment 2, that the target is (actually) not less likely to be located at the frequent distractor position as at any of the rare locations. In this situation, switching to dimension-based inhibition is adaptive: it minimizes distractor interference while not harming target processing at the frequent location.

Concerning question 2, it appears that observers adapt their mode of suppression to the prevailing positional distractor *and* target probabilities. When one location is highly likely to contain a distractor, as in Wang and Theeuwes’s (2018a) paradigm, the default set appears to be priority-map-based, which immediately brings about a strong reduction of distractor interference; a shift to dimension-based suppression is set in motion only later, when it is realized (over the course of the first session) that this set is associated with a substantial cost in processing targets at the frequent location. In contrast, in the paradigm of Sauter et al. (2018a), the distractor occurs randomly at various locations within a larger frequent (vs. a rare) distractor region, so that specific distractor positions are less obvious. In this case, the default may be dimension-based suppression, as the distractor is consistently defined – and, in fact, by the same feature – in a non-target dimension. The random swapping of the (distractor, non-distractor) color assignment across trials in the Wang and Theeuwes (2018a) paradigm (in contrast to the consistent assignment in the Sauter et al., 2018a, paradigm) may also retard adoption of a dimension-based set, as observers could not tell by the color of a stimulus that it is likely a distractor, rather than a target (but they could tell this more reliably based on its position at the frequent location). Thus, if spatial information is perceptually dominant over dimensional (or featural) information, observers may (first) come to operate a purely spatial (priority-map-based) distractor inhibition set.

More generally, this is to say that different default sets, or strategies, may be suggested by specifics of the individual paradigms, and overcoming these default sets may take time and additional learning of more subtle (e.g., target location) probability cues entailed in these paradigms. Thus, in a sense, the mode of suppression applied is in principle flexible. This does not necessarily mean that adopting a specific set or changing set involves a conscious decision; rather, it may simply be an adaptive process, driven by the availability of various distractor‐ and (target-) related probability cues. Also, it is conceivable that the two sets do not operate in an all-or-nothing fashion; rather, (as suggested by the roughly halved, though still significant target-location effect in the second compared to the first half of session 1 of Experiment 2) priority-map-based suppression may coexist with dimension-based suppression. However, more work is necessary to examine how this ‘mixture’ comes about: do the two sets operate in parallel within a given trial, or can only one set be effective on a trial (yielding a statistical mixture of the two sets across trials)?

In any case, the ‘locus’ of inhibition is flexible: priority-map‐ or dimension-based. And: just because one finds a distractor-location effect, one cannot conclude from this finding alone that inhibition operates at the level of the priority map. Ultimately, of course, it is the priority map via which the inhibition is always expressed, but the true level, at which it is instantiated at least in certain conditions, may be below the priority map. This is as envisaged by the dimension-weighting account, according to which selection is based on the priority map which, however, is itself shaped by the weighting applied to the various, target‐ and distractor-defining feature dimensions (for a recent review, see Liesefeld, Liesefeld, Pollmann, & Müller, 2018).

Finally, a few remarks are in order concerning other influences in the present paradigm, in particular, inter-trial effects.

The first concerns *positional cross-trial inhibition of distractor locations*. We did find evidence of passive carry-over of inhibition, however the effect tended to be small and relatively larger for rare distractor locations. The fact that there was hardly any effect for the frequent location lends support to the argument that distractors at this location are effectively suppressed by other means, limiting the room for passive cross-trial inhibitory effects to influence performance (if a distractor at this location is prevented from capturing attention, it does not need to be ‘extra’ suppressed, so no extra inhibition is carried over across trial). Thus, contrary to our initial hunch, positional cross-trial inhibition has next to no influence in the present paradigm – which may not be entirely surprising given that these effects tend to be small in any case with cross-dimensionally defined targets and distractors (see Sauter et al., 2018a, who found these effects to be larger by a factor of 4 with distractors defined in the same vs. a different dimension to the target).

The second concerns *color-based cross-trial repetition effects*. We did find *color-based repetition effects* (see Figure 7, which presents the effect pattern combined across Experiments 1 and 2), which were however relatively weak and tended to reflect, in the main, carry-over of (inhibition) of the distractor color from the previous trial, which aids performance if a same-colored distractor is present on the current trial (while it makes little difference with regard to where the current distractor appears, at the frequent vs. the rare locations). This contrasts with Wang and Theeuwes (2018a), who probably did not have the power to resolve these effects. In any case, the fact that there is a significant color-repetition benefit also for (current) trials with a distractor at the frequent location would indicate that the suppression of the distractor at this location is not entirely space-based, but includes some element of (albeit spatially parallel) color-based suppression. However, given that such an effect is seen only in the relatively infrequent event that there is a distractor at a rare location on the preceding trial (i.e., *p* = 0.66 • 0.35 • 0.66 • 0.65 ≈ 0.1, and *p* = 0.05 for color-repetition trials), and also given that the effect is equally seen when the distractor on trial *n* occurs at a rare location, carry-over of inhibition of the distractor color (from trial *n-1* into trial *n*) cannot account for (or can account at best for a small fraction of) the total interference reduction with distractors at the frequent versus one of the rare locations. This is the reason why we also find no difference (or only a numerical difference) when we compare simply the interference reduction for the frequent versus the rare locations between trials with a color repetition versus a switch (from the preceding trial): Experiment 1, 78 versus 67 ms, *t*(23)=1.27, *p*>0.05, *BF*=0.44; Experiment 2, 70 versus 83 ms, *t*(23)=−1.34, *p*>0.05, *BF*=0.48.

**Figure 7.**
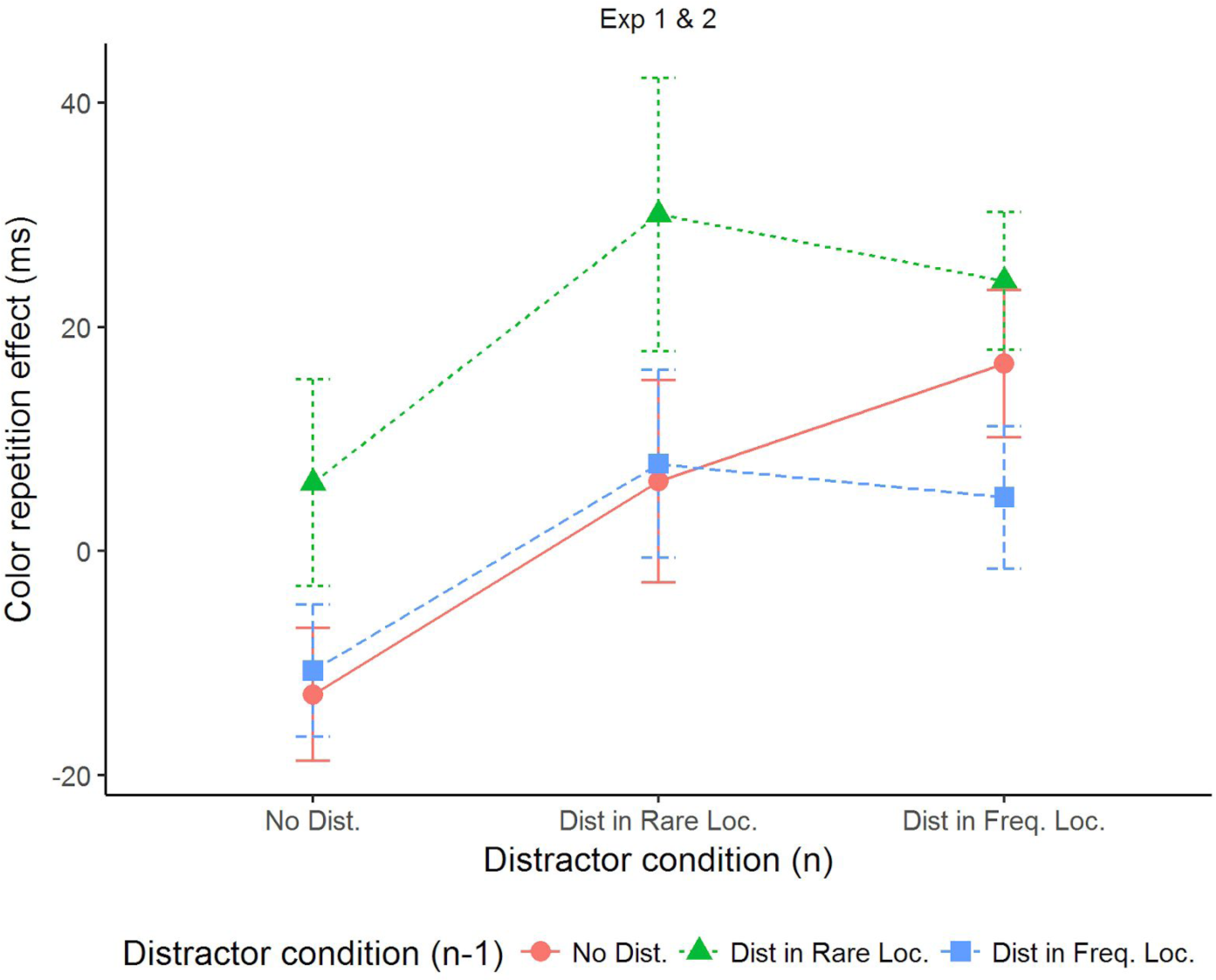
*Color-repetition effects (mean difference in RT between trials with a cross-trial change vs. repetition of the color assignment) as a function of the distractor condition on trial n, dependent on the distractor condition on trial n-1, combined across Experiments 1 and 2. Positive values mean RTs were faster when the same color was repeated (color-repetition benefits).*

With regard to the dimension-weighting account, one interesting issue in this context is why a color-feature-based modulation (spatially parallel carry-over of inhibition of the distractor color from the previous trial) appears to coexist with a dimension-based modulation (dimension-based suppression of the likely distractor location) in the second session of Experiment 2. As argued elsewhere (see, e.g., Sauter et al., 2018a), the two may not be incompatible: one may de-prioritize (down-weight) some specific feature at a feature-based level, while also de-prioritizing (down-weighting) the respective feature dimension at a higher level, prior to the integration of the dimension-specific feature contrast signals into the search-guiding (feature-less and supra-dimensional) priority map.

In sum, our finding (in two experiments) of a feature-based component of distractor suppression provides evidence in favor of all three levels – the featural, dimensional, and priority-map level (see Gaspelin & Luck, 2018, for a similar distinction) – being of importance.

A final word concerns the notion of an inhibitory gradient centered on the frequent distractor location. While we found some evidence of a gradient of inhibition on distractor-present trials in Experiment 1 (consistent with Wang & Theeuwes, 2018a), there was little evidence of a gradient effect on distractor-absent trials (not formally analyzed by Wang and Theeuwes) or distractor-present and ‐absent trials in Experiment 2 – despite the fact that our distractor location effects (the difference in RTs between frequent and rare locations) were comparable in magnitude to those reported by Wang and Theeuwes (2018a). Thus, it might be debatable whether the suppression effect is tightly centered on the frequent distractor location, or more fuzzily distributed around this location. The fact that there was no evidence whatsoever of a gradient effect in Experiment 2 might also suggest that the spatial extent of inhibition is influenced not only by the probability of the distractor occurring at the frequent location, but also by that of the target appearing in its vicinity. This requires more systematic investigation.

## CONCLUSION

We conclude that in the Wang and Theeuwes’s (2018a; see also Wang & Theeuwes, 2018b) paradigm, the learnt distractor location inhibition is not necessarily based on the priority map (as assumed by Wang & Theeuwes, 2018a; see also Ferrante et al., 2018), but may be shifted to a lower, dimension-based level in the functional architecture of search guidance – especially when the target location probability ‘affords’ a dimension-based inhibitory set, as this would leave target processing unaffected in scenarios in which the distractor is defined in a different dimension to the target (cf. Sauter et al., 2018a). On the other hand, observers do not mandatorily operate a dimension-based suppression strategy with different-dimension distractors (as implicitly assumed by Sauter et al., 2018a). Rather, at least when a single location is highly likely to contain a distractor, suppression at the level of the priority map may provide a ready ‘default’ strategy which is only slowly adapted in response to more subtle target location probability cues. Thus, both strategies are feasible in principle, and which one is adopted depends on the various, distractor and target probability cues acquired over the course of practice on the task.

## Acknowledgement

This work was supported by German Research Foundation (DFG) grants MU773/16-1, awarded to HJM and ZS, and MU773/14-1, awarded to HJM, as well as a China Scholarship Council (CSC) award to BZ. Correspondence concerning this article should be addressed to Hermann J. Müller, General and Experimental Psychology, Ludwig-Maximilians-Universität München, 80802, Munich, Germany.

## SUPPLEMENTARY

### Color-Repetition Benefits

Figure S1 depicts the RT benefit for repeated versus switched color assignments averaged across sessions, more precisely: the color repetition benefit as a function of the distractor condition (distractor absent, at rare location, at frequent location) on the current trial *n*, dependent on the distractor condition of *trial n-1*, for Experiments 1 and 2 (left and right panels), respectively.

**Figure S1.**
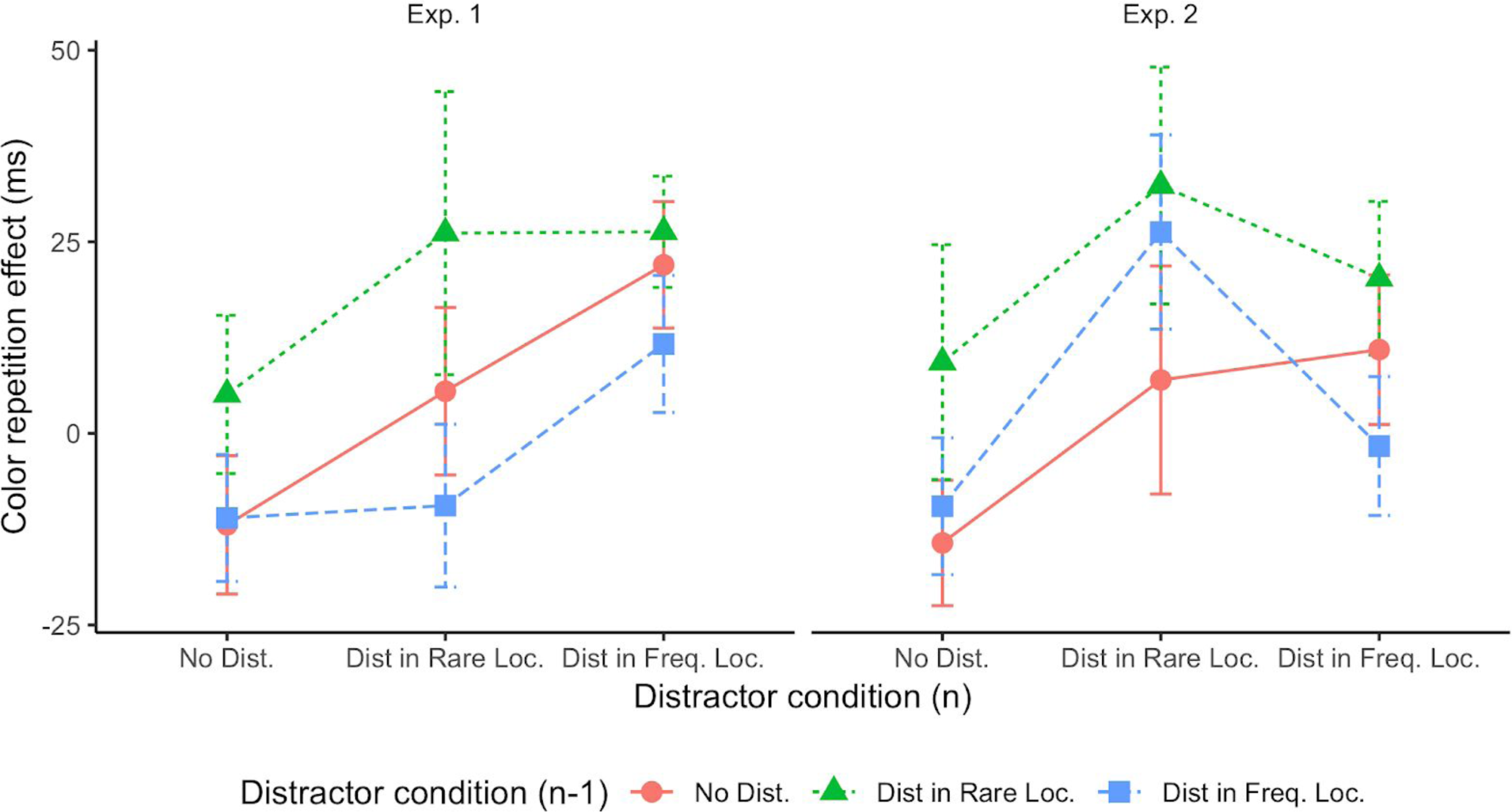
*The effect on RTs of repeating minus switching the target/distractor color between consecutive trials as a function of the distractor condition on trial n, dependent on the distractor condition on trial n-1. Positive values indicate that RTs were faster when the color was repeated.*

### Color-repetition effects in Experiment 1

As can be seen from the left panel of Figure S1, there was an RT benefit of repeating (vs. changing) the color assignment (distractor and non-distractor colors) from the previous trial when a distractor was present (vs. absent) on the current trial *n*, and this color repetition benefit was most marked when there was a distractor on the preceding trial *n-1*. A repeated-measures ANOVA revealed both main effects to be significant: distractor condition on trial *n* (*F*(38.8, 1.7)=4.01, *p*=0.032, 
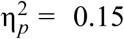
, *BF* = 2.5, Hyunh-Feldt corrected degrees of freedom) and distractor condition on trial *n-1* (*F*(46,2)=5.28, *p*=0.009, 
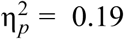
, *BF*=3.1); the interaction was not significant (*F*(70.3, 3.1) = 0.40, *p*=0.76, *BF*=0.057, Hyunh-Feldt corrected degrees of freedom). For interpreting the data, though, it should be borne in mind that some of the (cross-participant) condition means are likely to be beset by noise because the individual participants’ estimates (means) are based on very few observations (e.g., trials on which a distractor occurred at a rare location twice in a row (same or different rare location) were relatively infrequent, *p*=.054, i.e., *p*=0.027 for color repetition and switch trials). Overall, it appears that a (positive) color repetition manifested consistently (only) on a given trial *n* when, on the preceding trial *n-1*, a distractor had occurred at a rare location (explaining the main effect of the distractor condition on trial n-1). This produced a benefit especially when a distractor was also present on the current trial *n* at the frequent location (26-ms benefit, *t*(23)=3.51, *p*=0.02 (Bonferroni-corrected for multiple comparisons), *BF*=20.3), though there was also a numerical effect, of near-equal magnitude (30 ms), when a distractor was present at a rare location (though this benefit was statistically non-reliable: *t*(23)=1.72, *p*>0.05 (Bonferroni-corrected), *BF*=0.77).

### Color-repetition effects in Experiment 2

Similar to Experiment 1, there was an RT benefit of repeating the color assignment from the previous trial when a distractor was present on the current trial *n*, and (judging from the right panel of Figure S1) this color repetition benefit appeared to be most marked when there was a distractor on the preceding trial *n-1*. However, a repeated-measures ANOVA of the color repetition effect, with the factors distractor condition on trial *n* and distractor condition on trial *n-1*, revealed only the main effect of distractor condition on trial *n* to be significant (*F*(46,2)=5.78, *p*=0.006, 
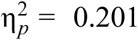
, *BF*=6.7), while the effect of distractor condition on trial *n-1* failed to reach significance: *F*(46,2)=2.00, *p*=0.15, *BF*=0.34) (interaction: *F*(63.1, 2.7)=0.41, *p*=0.80, *BF*=0.056). The significant main effect of distractor condition on trial *n* was due to the color repetition benefit being significantly larger than zero only when the distractor occurred at one of the rare locations (23 ms, *t*(23)=2.61, *p*=0.047 (Bonferroni corrected), *BF*=3.3), but not when it occurred at the frequent locations (12 ms, *t*(23)=1.67, *p*>0.1, *BF*=0.72). Note that when no distractor was present on trial *n*, there was not even a numerical benefit (if anything, there was a numerical, 7-ms cost; *t*(23)= −1.17, *p*=0.25, *BF*=0.39), consistent with Experiment 1 (5-ms cost; *t*(23)=−1.10, *p*=0.28, *BF*=0.37). Although this pattern looks the other way round to that seen in Experiment 1 (where the benefit was significant for the frequent location, but not for the rare locations), it should not be over-interpreted given the noise in the data (see above).

Thus, a possible account for the pattern common to both experiments (see also Figure 7 in the main text, which presents the data combined across the two experiments) may be as follows: When a distractor at a rare location captures attention (which is more likely to occur in comparison with a distractor at the frequent, i.e., ‘spatially’ suppressed, location), the distractor color is inhibited (and perhaps the non-distractor color enhanced) in order to disengage attention from the rare distractor and re-allocate it to the target. If this color set (inhibition of the distractor color, and perhaps facilitation of the target color) is carried over across trials, it would diminish the potential of a distractor defined by the same color, wherever it appears in the display, to attract attention (and a positive bias for the non-distractor color would help guide attention towards the target). Assuming a positive bias towards the non-distractor color (in addition to a negative bias towards the distractor color) would explain the slight numerical benefit seen even if there is no distractor present on the current trial. Also in line with this account is the fact there is no significant color-repetition benefit with a distractor at the frequent location on trial *n-1*: as such distractors (at the spatially suppressed location) are unlikely to capture attention, their color is not encoded (and inhibited), thus not giving rise to a color repetition effect. – This, arguably, makes sense of key features of the pattern seen in Figure S1, although this pattern appears to be richer than the post-hoc account sketched here. Further, dedicated work would be required to explore these more subtle effects.

## Footnotes

1 Sauter et al. (2018a) took the finding that, with same-dimension distractors, the reduction of distractor interference was accompanied by impaired target processing to argue against distractor suppression operating at a feature-based level: if the distractor-defining feature could be selectively inhibited in the frequent distractor region, target processing should have been *un*impaired in this region not only when the distractor was defined in a different dimension to the target, but also when it was defined in the same dimension (in which case the two were still categorically separable, i.e., affording feature-based search guidance, in terms of Wolfe and Horowitz (2017): the target was a bar differing by a ±12° tilt from the vertical non-target bars, whereas the distractor was a horizontal, 90°-tilted bar).

2 We conducted directed *t*-tests as, based on previous studies (e.g., Sauter et al., 2018), we had predicted a coincidence effect.

3 The reduction of the target-location effect is numerically similar when removing trials from the analysis on which the target position on trial *n* coincided with the distractor position on trial *n-1* (second vs. first half: 62 vs. 100 ms), but not statistically significant (*t*(23)=1.44, *p*=0.16, *BF*=0.53). Note though that estimating the carry-over of inhibition of the previous distractor location to the current target location is inherently more noisy (especially for the rare locations) when the estimates are based on only half the number of trials.

4 Ruthruff and Gaspelin (2018) proposed a similar, ‘spatial-filtering’ account, to explain the lack of interference caused by a salient onset ‘pre-cue’ stimulus presented at one of two invariable, i.e., known, non-target locations in a variant of the ‘contingent-capture’ paradigm (cf. Folk & Remington, 1996).

5 In Experiment 1, 12 (of the 24) participants correctly pointed to the likely distractor location in an eight-alternative forced-choice test at the end of the second session, and 14 participants if including those who indicated a position directly adjacent to the true location. In Experiment 2, 6 (of 24) participants correctly pointed to the likely location, and 13 when those are included who indicated a position adjacent to the true location. Thus, 56.25% of the participants had precise (37.5%) or approximate (18.75%) knowledge of the likely distractor location, which compares with a chance-expectancy level of 37.5% (i.e., 12.50% plus 25.00% for the precise plus the adjacent locations). This indicates that participants had above-chance, explicit knowledge of the likely distractor location. This is consistent with Wang and Theeuwes (2018a), who had found no difference in performance between observers who could or could not explicitly tell the frequent distractor location. That is, suppression of the likely distractor location reflects, by and large, an implicit learning effect (see also Sauter et al., 2018a).

